# Species variations in tenocytes’ response to inflammation require careful selection of animal models for tendon research

**DOI:** 10.1101/2021.05.09.443263

**Authors:** Gil Lola Oreff, Michele Fenu, Claus Vogl, Iris Ribitsch, Florien Jenner

## Abstract

For research on tendon injury, many different animal models are utilized; however, the extent to which these species simulate the clinical condition and disease pathophysiology has not yet been critically evaluated. Considering the importance of inflammation in tendon disease, this study compared the cellular and molecular features of inflammation in tenocytes of humans and four common model species (mouse, rat, sheep, and horse). While mouse and rat tenocytes most closely equalled human tenocytes’ low proliferation capacity and the negligible effect of inflammation on proliferation, the wound closure speed of humans was best approximated by rats and horses. The overall gene expression of human tenocytes was most similar to mice under healthy, to horses under transient and to sheep under constant inflammatory conditions. Humans were best matched by mice and horses in their tendon marker and collagen expression, by horses in extracellular matrix remodelling genes, and by rats in inflammatory mediators. As no single animal model perfectly replicates the clinical condition and sufficiently emulates human tenocytes, fit-for-purpose selection of the model species for each specific research question and combination of data from multiple species will be essential to optimize translational predictive validity.

## Introduction

Animal models are cornerstones of biomedical and translational medicine research. They are used when it is unethical or impractical to study the target species to explore basic pathophysiological mechanisms, to evaluate safety and efficacy of new treatment approaches, and to decide whether novel therapeutic candidates warrant the economic and moral costs of clinical development.^1-7^ For 90% of new treatment strategies, however, translation from basic science to the clinic fails, mainly because clinical trials show them to be inefficient (52%) or unsafe (24%) during phases II and III.^4,5,8^ Such translational failures cost animal lives, strain clinical trial volunteers, and burden biomedical research, the pharmaceutical industry and health care systems. So far, attempts to optimize translational success have mainly focused on internal validity flaws such as methodological shortcomings in animal and clinical trials, publication bias, or overoptimistic conclusions about efficacy. Yet another key factor, the external validity, or generalizability, of animal models has received little attention.^4,5,8-13^ Common problems of external validity include species differences in disease pathophysiology, common confounding comorbidities and the selection of outcome measures.^5^ An animal model should sufficiently emulate aetiology, pathophysiology, symptomatology and response to therapeutic interventions of the target species to allow extrapolation.^5,11^ As no single animal model perfectly recapitulates the clinical realm, fit-for-purpose validation and selection of the most appropriate model species is essential.^10-13^ Unfortunately, for musculoskeletal disorders, such as tendinopathy, in depth validation studies of animal models beyond structural and biomechanical similarities are largely lacking.

Tendinopathy, a disabling overuse injury, is the most common musculoskeletal complaint for which patients seek medical attention.^14^ It is prevalent in both occupational and athletic settings, afflicting 25% of the adult population, and accounting for 30–50% of all sport injuries.^15-18^ Major tendons experiencing high loads are most commonly affected, especially the weight-bearing and energy-storing Achilles tendon, which routinely experiences loads of up to 12.5 times the weight of the individual.^6,19^ Many intrinsic and extrinsic factors, including age, body weight and physical loading, influence the aetiopathogenesis of tendinopathy. Overload and repetitive strain lead to accumulation of microdamage and concurrent inflammatory, dysregulated reparative and degenerative processes, causing clinical symptoms, e.g., activity-related pain, focal tendon tenderness and swelling, and functional limitations. Overt clinical symptoms such as pain are preceded by tendinous matrix remodelling, an inflammatory cellular process mediated in part by metalloproteinase enzymes.^20,21^ Due to its low cellularity, vascularity and metabolic rate, a tendon’s response to injury is inefficient, requiring lengthy periods of recuperation and often resulting in a fibrovascular scar. Scar tissue has significantly inferior biomechanical properties than the original tendon tissue and is prone to re-injury.^16,22-24^ Current treatment options are mostly palliative and fail to restore the functional properties of injured tendons.^16,22-24^ Tendinopathy thus has a significant adverse impact on quality of life and costs individuals and the society an estimated annual expense of $30 billion.^25,26^ This is driving research efforts into unravelling the molecular mechanisms of tendinopathy and developing targeted regenerative therapies. Of particular interest in this context are the cellular and molecular processes orchestrating inflammation in tendinopathy and the mechanisms governing the development of chronic inflammation that fails to resolve in persistently symptomatic patients.^27^

Tendon injury induces a local inflammatory response, characterized by immune cell infiltration and the expression of pro-inflammatory mediators, which in turn reduce collagen production and induce vasodilation, angiogenesis, and matrix metalloproteinase (MMPs) expression.^28-31^ Furthermore, the inflammatory milieu can modify tenocyte physiology by increasing metabolic activity and inducing an activated, proinflammatory phenotype with inflammation memory and the capacity for endogenous production of cytokines such as TNF-α, IL-1β, IL-6, IL-10, VEGF and TGF-β.^29,30,32^ While the initial inflammatory response is essential to start the healing process, sustained inflammatory conditions contribute to dysregulated matrix remodelling and fibrovascular scarring during healing.^18,31^ Chronic inflammation thus drives tendon degeneration before tearing or any other clinical signs of tendinopathy, impairs healing after injury and promotes the development of tendinopathy.^14^

While human tendon tissue can typically be procured only from individuals with advanced pathology, animal models provide the opportunity to obtain tissue during all stages of tendinopathy to study organ, cellular and molecular changes over the entire course of the disease. In animal models, consistent and repeatable injuries can be induced, evaluated and treated, while controlling for potential confounding influences.^23,33,34^ Since no species has been established as the gold standard for tendinopathy research, many induced and spontaneous animal models ranging from small rodents (mice, rats) to large animals (sheep, horses) are utilized.^19,35-43^ While the biomechanical properties of the various species are well established, their ability to simulate the pathophysiology of human tendon disease, including the molecular behaviour of key genes and pathways, has not been critically evaluated yet and detailed analyses of species-specific differences in cytokine expression and regulation as well as of tenocytes susceptibility to cytokines are still lacking.^44^

Considering the importance of inflammation in tendon disease,^29,33,45,46^ this study compares the cellular and molecular features of inflammation in tenocytes of humans and four common model species (mouse, rat, sheep, and horse) to aid in the evidence-based selection of fit-for-purpose translational animal models for tendon research. Mice and rats are included due to their prevalent use as laboratory animals and availability of species-specific molecular tools.^6,47^ Larger animals are used increasingly as translational models due to their more comparable tendon dimensions and biomechanics.^23,48-50^ Horses present an attractive model of human tendinopathy since their superficial digital flexor tendon is a weight-bearing and energy-storing tendon analogous to the human Achilles tendon, which is similarly prone to naturally occuring tendon disease with high recurrence rates.^9,51,52^ Furthermore equine ageing proceeds similarly to humans.^39,51,52^ Sheep are included because features of clinical tendinopathy of horses could be emulated also in ovine induced models.^51,53-57^ In particular, the ovine intra-synovial tendon lesion model mimics the clinical intra-synovial tendon disease of humans and horses more accurately than small animal extra-synovial models, e.g., with respect to histology and gene expression, to similarities in the biomechanical environment and to failure of lesions to heal.^51,53-57^

## Results

### Morphology

Tenocytes from all five species shared common characteristics with a fusiform appearance, adherence to the flask and similar dimensions (fig. 1): human tenocytes measured 177.8 ± 40.1 μm (mean ± s.d.) in length and 20.2 ± 4.4 μm in width, mouse 163.7 μm (±23.6) x 19.4 μm (±3.3), rat 182.5 (±26.6) x 24.2 (±8.0), sheep 206.3 (±45.5) x 28.9 (±8.5) and horse 193.2 (±8.4) x 15.4 (±1.4). In high confluency, tenocytes from human, sheep and horse showed similar morphology, creating cell bundles arranged in a storiform pattern (fig. 1A), while tenocytes from mouse and rat had a more scattered appearance with a random orientation.

**Figure 1:**
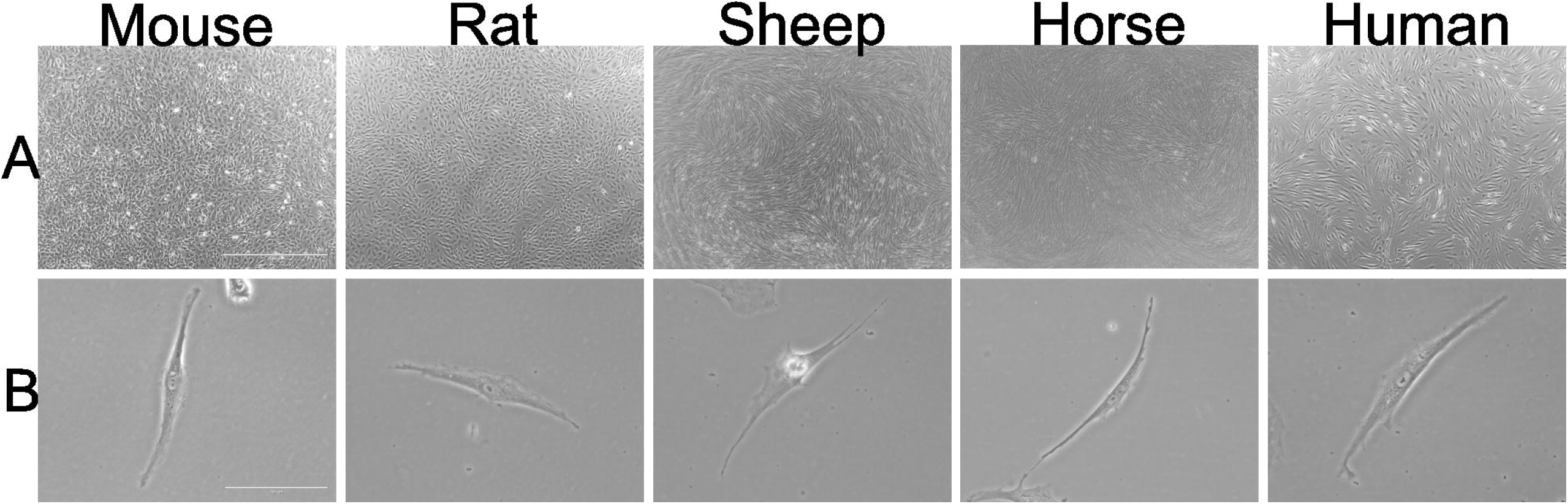
Micrographs of tenocytes derived from the Achilles tendon in the mouse, rat, sheep and human or the superficial digital flexor tendon in the horse. Figure 1A shows the tenocytes at a magnification of 40x (scale bar: 1000μm), while figure 1B was taken at a 400x magnification (scale bar: 100μm).

### Proliferation assay

Proliferation, migration and gene expression of tenocytes of all five species were compared under standard culture conditions (healthy control) as well as under transient (24 h) and constant exposure to inflammatory stimuli (fig. 2). Under healthy conditions (fig. 3, table 1), equine tenocytes had the highest proliferation rates while murine cells had the lowest. Human tenocytes exhibited the second lowest proliferation capacity with an inability to double the cell amount over the 48h observation period. Sheep and rat tenocytes were in the middle. From fig. 3, it can be seen that slopes are quite variable among species, but within species variation is low. Therefore, the slopes of tendon cells of all four model species are significantly different from those of humans (p<0.001; to correct for multiple testing using Bonferroni with four comparisons, the nominal significance levels, 0.05, 0.01, and 0.001, are set to the corrected levels: 0.0125, 0.002, 0.0002, respectively), even for only three biological replicates per species. Under constant exposure to inflammatory stimulation (10 ng/ml IL1β and 10 ng/ml TNFα), the proliferation of sheep tenocytes decreased significantly and fell to about human levels, i.e., the difference in the proliferation slopes between healthy and constantly inflamed sheep decreased significantly (p<0.01). All other differences in slopes between healthy, constantly and transiently (only 24h inflammatory stimulation) inflamed conditions were not significant.

**Figure 2:**
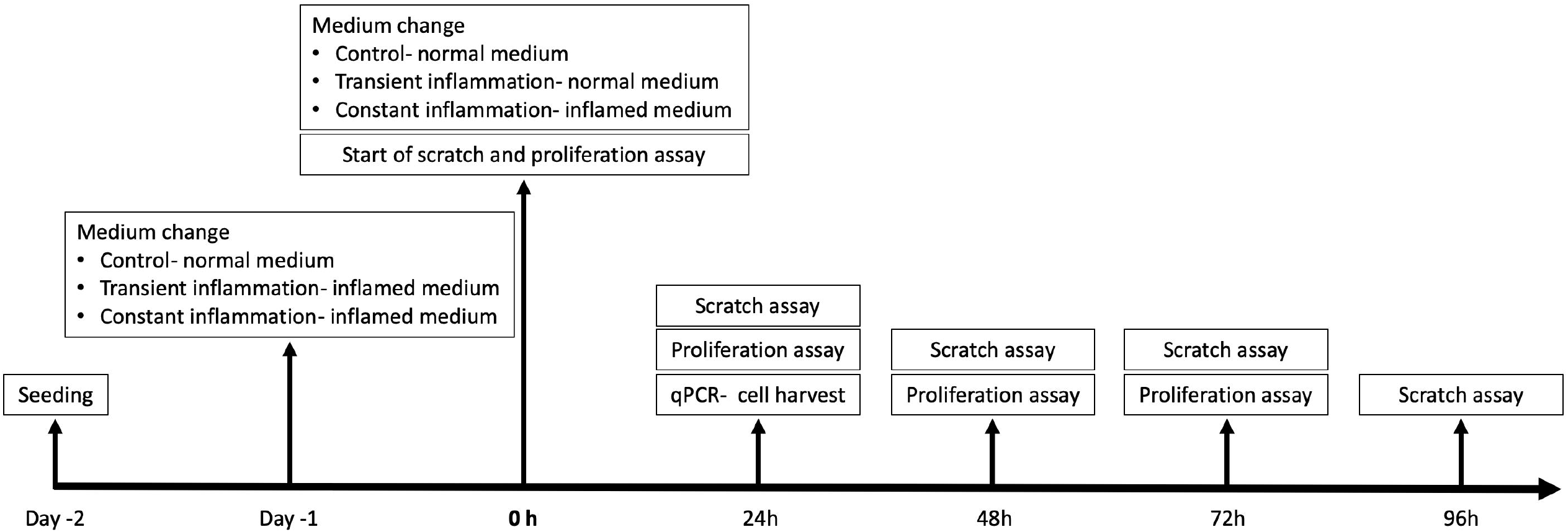
Study timeline detailing the experimental protocol. Gene expression, proliferation and migration of tenocytes of all five species (human, mouse, rat, sheep, horse) were compared under standard culture conditions (healthy control) as well as under transient (24 h) and constant exposure to inflammatory stimuli (10 ng/ml IL1β and 10 ng/ml TNFα). After 24-hour exposure to inflammation, the transient inflammation group received fresh culture medium, while for the constant inflammation group fresh medium was again supplemented with inflammatory factors.

**Figure 3:**
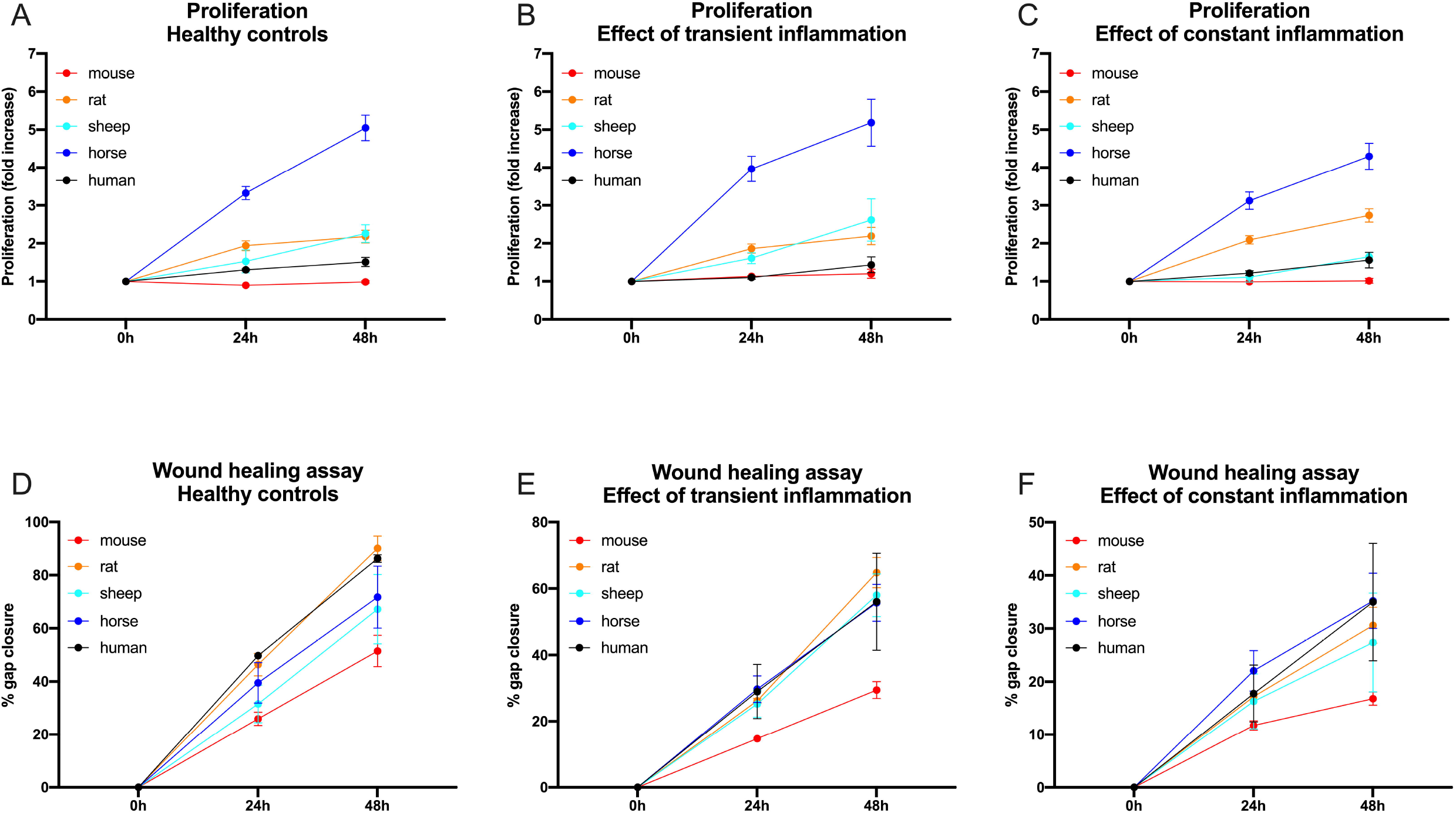
The proliferation capacity (A-C) of tenocytes from 5 different mammalian species (mouse, rat, sheep, horse, and human) under healthy (Ctr/A), transient (TI/B) and constant (CI/C) inflammatory condition is illustrated as fold increase over the course of 2 days (indicated as mean ± SEM calculated from three biological replicates). For pairwise comparisons and significance values see table 1. A wound healing assay (D-F) was used to determine the migratory capacity of tenocytes from five different mammalian species under healthy (D), transient (E) and constant (F) inflammatory conditions (indicated as mean ± SEM calculated from three biological replicates). For pairwise comparisons and significance values see table 1.

**Table 1:** The first column corresponds to the healthy situation, the second and third to transiently and constantly inflamed, respectively. First rows (human) correspond to tests for non-zero slopes of the proliferation and migration curves in humans, i.e., either non-zero proliferation or non-zero gap closure in humans. Second to fifth rows (different animal species) correspond to tests of differences of the animal models to humans in the slopes of the proliferation and migration curves of tenocytes. Calculations used an ANCOVA. Means, standard errors and p-values are reported. To correct for multiple testing with four comparisons using Bonferroni, the nominal significance levels (0.05, 0.01, and 0.001) are set to the corrected levels (0.0125, 0.002, 0.0002, respectively). P-values are marked with stars from * (significant) to *** (highly significant) using this correction.

| SPECIES |  | HEALTHY |  |  | TRANSIENT INFLAMMATION |  |  | CONSTANT INFLAMMATION |  |  |
| --- | --- | --- | --- | --- | --- | --- | --- | --- | --- | --- |
|  |  | *prolif. slope* | *Std. error* | *p-value* | *slope: diff. to healthy* | *Std. error* | *p-value* | *slope: diff. to healthy* | *Std. error* | *p-value* |
| PROLIFERATION | Human | 0.1027 | 0.0168 | 1.75e-09 *** | -0.0167 | 0.0237 | 0.4801 | 0.0089 | 0.0237 | 0.7074 |
|  |  | *slope: diff. to human* | *St. error* | *p-value* | *slope: diff. to human* | *St. error* | *p-value* | *slope: diff. to human* | *St. error* | *p-value* |
|  | Mouse | -0.1250 | 0.0237 | 1.95e-07 *** | 0.0508 | 0.0335 | 0.1301 | -0.0301 | 0.0335 | 0.3691 |
|  | Rat | 0.0933 | 0.0237 | 9.35e-05 *** | -0.0038 | 0.0335 | 0.9103 | 0.0420 | 0.0335 | 0.2103 |
|  | Sheep | 0.1169 | 0.0237 | 1.10e-06 *** | 0.0585 | 0.0335 | 0.0815 | -0.0878 | 0.0335 | 0.00900 * |
|  | Horse | 0.3000 | 0.0237 | < 2e-16 *** | 0.0233 | 0.0335 | 0.4868 | -0.0640 | 0.0335 | 0.0568 |
|  |  | *migration slope* | *Std. Error* | *p-value* | *slope: diff. to healthy* | *St. error* | *p-value* | *slope: diff. to healthy* | *St. error* | *p-value* |
| MIGRATION | Human | -<br>0.0792257 | 0.0032 | < 2e-16 *** | 0.0277 | 0.0045 | 2.76e-09 *** | 0.0475 | 0.0045 | < 2e-16 *** |
|  |  | *slope: diff. to human* | *Std. Error* | *p-value* | *slope: diff. to human* | *St. error* | *p-value* | *slope: diff. to human* | *St. error* | *p-value* |
|  | Mouse | 0.0311503 | 0.0045 | 3.07e-11 *** | -0.0075 | 0.0064 | 0.2452 | -0.0156 | 0.0064 | 0.0154 |
|  | Rat | 0.0001854 | 0.0046 | 0.9155 | -0.0063 | 0.0064 | 0.3287 | 0.0035 | 0.0065 | 0.5938 |
|  | Sheep | 0.0185550 | 0.0045 | 5.46e-05 *** | -0.0188 | 0.0064 | 0.00374 * | -0.0112 | 0.0064 | 0.0811 |
|  | Horse | 0.0132563 | 0.0045 | 0.00375 ** | -0.0123 | 0.0064 | 0.0567 | -0.0140 | 0.0064 | 0.0302 |

### Wound healing (scratch) assay

Gap closure was significantly different (in all cases p<0.001) between all conditions, fastest for healthy and slowest for constantly inflamed cells of all species (fig. 3, table 1). Murine tenocytes showed the slowest wound healing for all conditions. There was no statistically significant difference in migration speed between healthy and transient inflammatory conditions in tenocytes of any species and between healthy and continuously inflamed conditions only for mouse and rat cells (p< 0.05).

Under healthy conditions, gap closure of tenocytes from all species except rats was significantly different (in all cases p< 0.003) from humans with cells from rats showing the fastest wound healing (gap closure at 48h mean 90.08% ± 8.01% s.d.), closely followed by humans (gap closure at 48h mean 86.25% ± 2.47% s.d.) and cells from mice the slowest (gap closure at 48h mean 51.49% ± 10.23% s.d.). Under transient inflammation rat tenocytes again were fastest (gap closure at 48h mean 64.81% ± 7.84% s.d.), with ovine (gap closure at 48h mean 58% ± 11.17% s.d.), human (gap closure at 48h mean 56.06% ± 25.27% s.d.) and equine tenocytes (gap closure at 48h mean 55.71% ± 9.6% s.d.) following with similar wound healing rates, while murine tendon cells again were slowest (gap closure at 48h mean 29.46% ± 4.36% s.d.). Under constant inflammation, equine tenocytes were fastest (gap closure at 48h mean 35.23% ± 8.98% s.d.) followed closely by human tendon cells (gap closure at 48h mean 34.99% ± 19.12% s.d.). The change in migration speed compared to healthy was significantly different from human tenocytes for ovine tendon cells under transient and equine and murine tenocytes under constant inflammation (table 1).

### Quantitative PCR

The species show variable approximations of human expression levels among functional gene groups and conditions (table 2 and 3, fig. 4 and 5, suppl. fig. 1 and 2, suppl. table 1).^58^ A univariate Analysis of Variance (ANOVA) demonstrated significant differences between each species and humans in many genes relevant for tendon function and inflammatory response (table 2). Remarkably, healthy tenocytes of all four species show significant differences to humans in their expression of Col1, the main tendon matrix component, of collagenase MMP13 and of the key inflammatory mediator COX2. Similarly, significant differences from humans in the COX2 expression of transiently inflamed tenocytes are evident for all four species and in IL6 expression for all species except rats. In contrast, no significant differences to humans were seen in MMP1 expression for any species and the osteogenic marker ALP was only significantly different in healthy murine tenocytes. NFkB expression exhibited a significant difference only in healthy horse tendon cells and p53 in healthy horse and rat tenocytes.

**Table 2:**
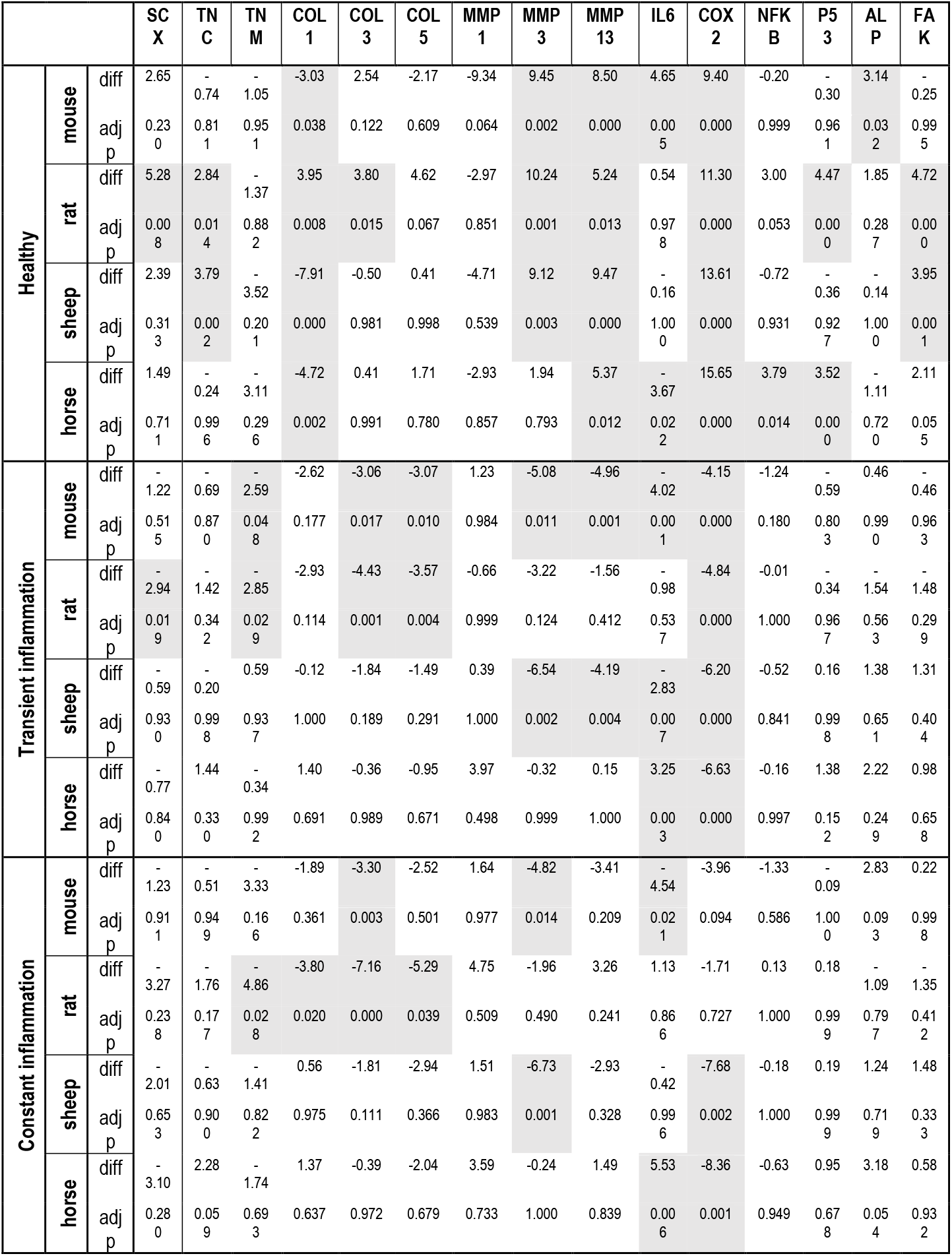
Mean difference to human and p-values of the gene expression calculated with ANOVA of healthy, transiently inflamed and constantly inflamed tenocytes of the four animal model species. Significant p-values (Tukey HSD correction) and the matching mean differences in gene expression to humans are indicated in bold.

**Table 3:**
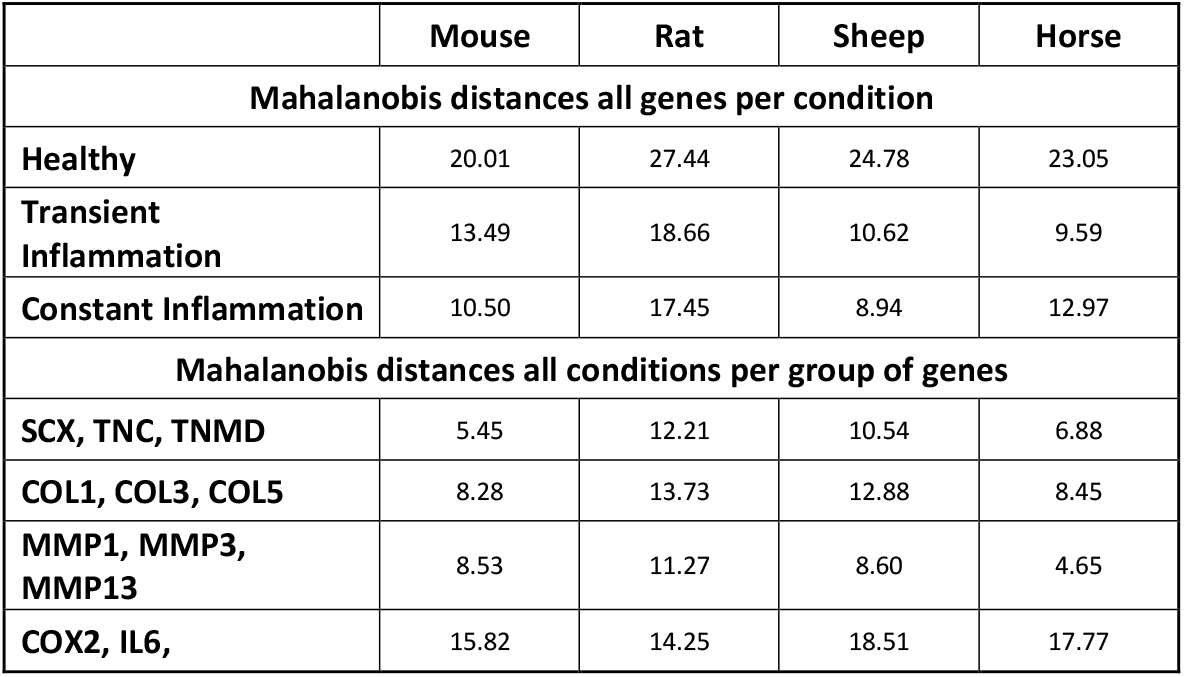
Mahalanobis distances of the four model species to humans for all genes combined under healthy, transiently and constantly inflamed conditions as well as for the different functional gene groups: tenogenic markers, collagens, MMPS and inflammatory mediators.

|  | Mouse | Rat | Sheep | Horse |
| --- | --- | --- | --- | --- |
| **Mahalanobis distances all genes per condition** |  |  |  |  |
| **Healthy** | 20.01 | 27.44 | 24.78 | 23.05 |
| **Transient Inflammation** | 13.49 | 18.66 | 10.62 | 9.59 |
| **Constant Inflammation** | 10.50 | 17.45 | 8.94 | 12.97 |
| **Mahalanobis distances all conditions per group of genes** |  |  |  |  |
| **SCX, TNC, TNMD** | 5.45 | 12.21 | 10.54 | 6.88 |
| **COL1, COL3, COL5** | 8.28 | 13.73 | 12.88 | 8.45 |
| **MMP1, MMP3, MMP13** | 8.53 | 11.27 | 8.60 | 4.65 |
| **COX2, IL6,** | 15.82 | 14.25 | 18.51 | 17.77 |

**Figure 4:**
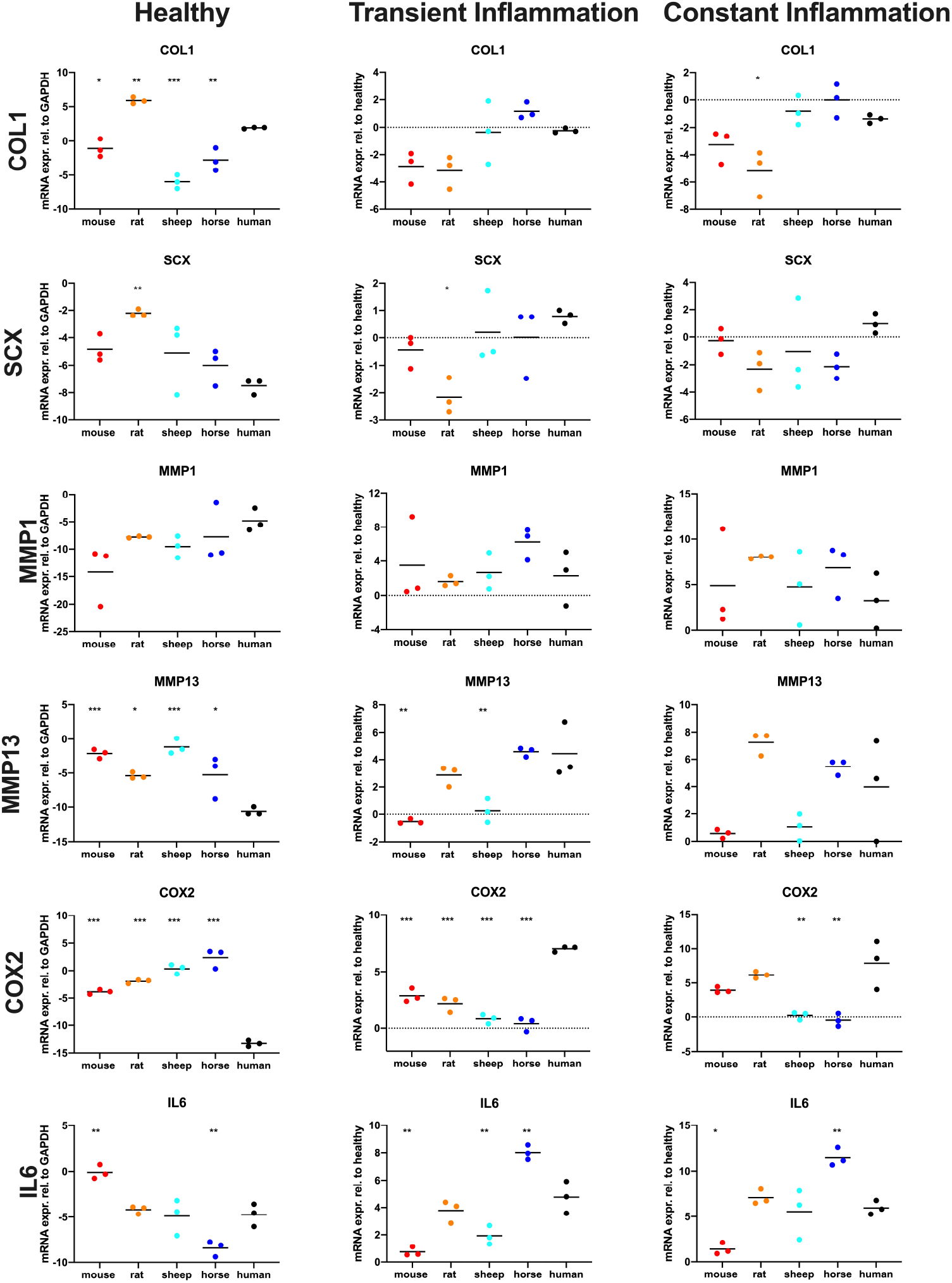
Scatter dot plots showing COL1, SCX, MMP1, MMP13, COX2 and IL6 gene expression (presented as log2) of healthy tenocytes and tenocytes exposed to inflammatory stimuli for 24h (transient inflammation) or continuously (constant inflammation) in different species (the black lines indicate the respective means). Gene expression in the inflammatory conditions is shown relative to the healthy tenocytes. Each dot represents a different biological replicate. Differences were evaluated using ANOVA with Tukey HSD test, *p<0,05; **p<0,01; ***p<0,001

**Figure 5:**
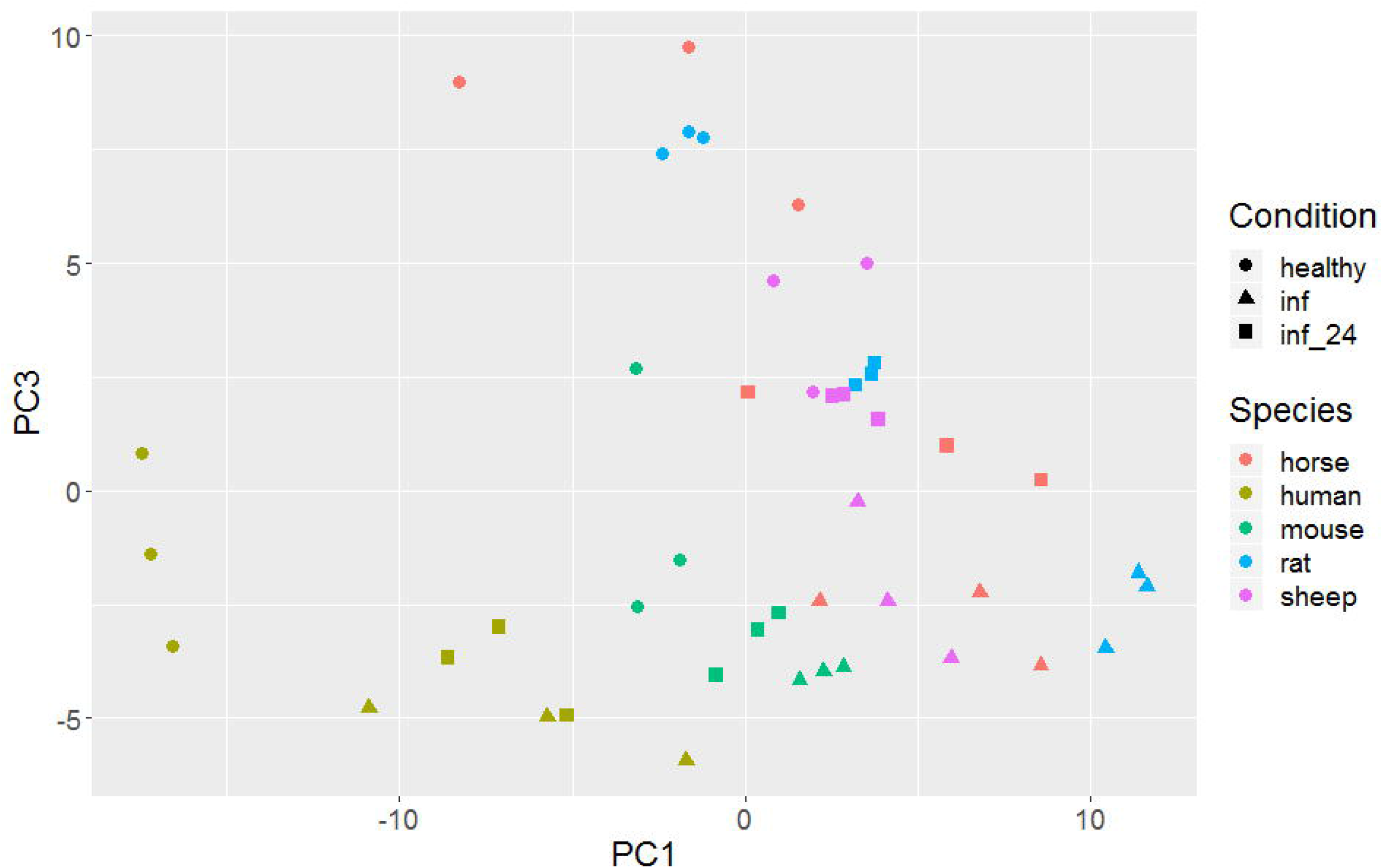
Plot of PC1 (explaining 36% of the variance) vs. PC3 (explaining 14% of the variance) of a Principal Component Analysis of gene expression values. Species are colour coded, conditions (healthy, transiently inflamed, and continuously inflamed) are differentiated by symbols. Each dot represents a different biological replicate.

To condense the information from the univariate ANOVA results to overall measures of similarity between the different animals and humans we calculated the multivariate Mahalanobis distances of the four species to humans. The Mahalanobis distance is a non-dimensional measure of dissimilarity, where between group distances are weighted by the inverses of within group variability, much like the test statistic of a t-test for one variable. Similar to its use in graphically detecting outliers in multiple dimensions, we use it to show multivariate dissimilarity in gene expression among species. In supplementary figure 1, we summarize conditions within a gene. From figure 4, it is evident that Cox2 is rather variable among species (especially human healthy cells differ from all animals, least so from mice) while showing relatively little variation within species under all conditions. Hence the Mahalanobis distances to humans are comparatively large with mice being closest to humans (suppl. fig. 1). The second immune gene, Il6, shows also relatively little within species variation (fig. 4), but comparatively less between species variation; especially rats and sheep are similar to humans, while mice and horses are slightly further away (fig. 4, suppl. fig. 1). The Mahalanobis distances (table 3) of the immune genes are relatively large compared to the other gene combinations due to low within species variability. The overall distance of the immune genes is a compromise between the two genes, such that rats are slightly more similar to humans than mice, with sheep and horses further away. When different functional groups of genes are analysed, variable Mahalanobis distances of the four model species from humans are found (table 3). While mice and horses are closest to humans in their tendon marker and collagen expression, horses appear to be by far the best model for MMPs (table 3). For the overall gene expression of healthy tenocytes, Mahalanobis distances from human tenocytes are large and similar among species, with mouse tendon cells appearing least and rat tenocytes most distant from human. For transient inflammation, Mahalanobis distances to humans are generally lower, but the spread of differences widens, with horses, sheep and mice relatively close and rats clearly furthest. For constant inflammation, the pattern is qualitatively similar to that under transient inflammation. Overall, the pattern of multivariate dissimilarity of species varies widely and unpredictably among species pairs with no single species most similar to humans.

Principal Component Analysis (PCA) is an exploratory technique for reducing the complexity of data. We used expression data from all genes and plot the results for the different species x condition combinations (fig. 5, suppl. fig. 2). The plot of PC1 vs. PC3 is easier to interpret than of other combinations of PCs. (fig 5). With the exception of humans and mice, species are generally not well-separated. Within all species, the different conditions are spread along an oblique line with healthy to the upper left and inflamed to the lower right. Within mice, healthy is also separated from inflamed, while in sheep the pattern is less clear. Pairwise plots of other PCs show similar but less clear patterns (suppl. fig. 2).

## Discussion

Choice of the most appropriate animal model is the most essential and challenging element of animal-based research, and also an important aspect of the 3Rs (i.e., replace, reduce, refine) to ensure the best use of animals.^59-61^ Unfortunately, the choice of animal models frequently is based more on convention, financial and practical considerations, such as housing and husbandry requirements or the availability of reagents and biochemical tests, than compelling scientific evidence of the fit to human diseases and clinical contexts.^4,5,9,62-64^ The lack of formal requirements for animal models is due to the traditional assumption that genetic homology derived from a common evolutionary origin also implies functional similarities of gene regulation, signalling pathways and developmental systems between species (the “unity in diversity” concept).^7^ Species may, however, differ in critical aspects and rarely have assumed similarities been empirically demonstrated.^7^ The diversity of human patients and symptoms is thus unlikely fully represented in highly inbred rodents.^65,66^ Even humanized models, which have contributed significantly to research by facilitating functional studies in vivo, cannot replicate the complexity of human disease.^67^

Both the European Medicines Agency (EMA) and the USA Federal Food and Drug Administration require the use of fit-for-purpose animal models to evaluate efficacy, durability, dose-response, degradation and safety of new therapeutics for market approval. Recently, these regulatory authorities published guidelines identifying requirements to demonstrate the relevance of animal models for investigational new product testing by cross-species comparison of the structural homology of the target, its distribution, signal transduction pathways and pharmacodynamics.^68,69^ Furthermore, several voluntary initiatives have established criteria to encourage the evidence-based selection of animal models for stroke and schizophrenia.^11,70,71^ To this end, both the model species and disease-induction protocols, need to be validated by comparing the animal model with the gold standard or the target species.^11^ As no gold standard for tendon research is available, this study compared tenocyte morphology, proliferation rate, wound healing speed, and gene expression of two small animals (mouse and rat) and two large animals (sheep and horse) to human under healthy as well as “diseased” (transiently and continuously inflamed) conditions to determine similarities and differences among species. It can serve as the foundation for a rational, evidence-based choice of optimal animal models for specific aspects of human tendinopathy.

Tendon injury induces a local inflammatory response, which initiates the healing process. Tendon healing occurs in three chronologic phases: inflammation (0-7 days), proliferation (1-6 weeks), and remodeling (6 weeks – 6 months). While these stages overlap, they are characterized by temporally and functionally distinct cytokine profiles and cellular processes.^72^ The initial inflammatory phase is characterized by influx of inflammatory cells, which release chemotactic and proinflammatory cytokines and growth factors that lead to recruitment and proliferation of macrophages and resident tendon fibroblasts.^44,72-83^ In addition, tenocytes produce also several endogenous cytokines and growth factors which contribute to the healing process in an auto- and paracrine manner.^44,75^ During the proliferative stage tenocytes proliferate and produce an immature neomatrix with a predominance of type III rather than type I collagen.^44,72,80-82,84^ Lastly, in the course of the remodeling phase, the cellularity decreases, matrix synthesis is reduced and collagen fibrils and tenocytes align linearly with the direction of tension.^44,72-79,81-83^ However, in both man and horse suffering from naturally occurring tendon disease, the normal architecture, composition and function of the tendon are never completely restored, predisposing them to recurring injury and tendinopathy.^44,72-79,81-83^

Given the importance of inflammation in tendon injury and repair, with pro-inflammatory cytokines acting as a regulatory link between several catabolic and anabolic systems and as a double-edged sword both promoting and impeding tendon repair,^44,83,85-88^ this study focused on the comparative response to inflammation.

We used IL-1β and TNF-α, two hallmark cytokines of inflammation in tendons, which are associated with tendon injury and tendinopathy in vivo and in vitro, to induce disease-relevant inflammation.^28,30,33,45,89-97^ IL-1 activates the NF-κB pathway in tenocytes, induces the production of inflammatory mediators including COX2 and IL6, and matrix remodelling factors such as MMP1, MMP3 and MMP13.^28,30,89-91^ It can even cause loss of the tenocyte phenotype, which is associated with decreased expression of tendon-related genes, e.g. COL1, SCX and TNMD.^28,30,89-91^ Similarly TNF-α can strongly activate tenocytes, stimulating them to produce more cytokines, including IL-1β, TNF-α, IL-6.^28,30,92-94,96^ Accordingly, we used tendon-specific markers (SCX, TNC, TNMD, COL1, COL3, COL5), matrix remodelling proteinases (MMP1, MMP3, MMP13) and inflammatory factors (COX2, IL6, NF-κB, p53) in addition to proliferation and wound healing speed as read-outs to evaluate the response to IL-1β/ TNF-α induced inflammation.

Interestingly, while mouse and rat tenocytes most closely matched human tenocytes’ low proliferation capacity and minimal effect of inflammation on proliferation, the human wound closure speed was best approximated by rats and horses. Tenocyte migration to the injured tissue and proliferation are essential processes in tendon healing.^98,99^ Accordingly, inflammatory stimulation, e.g. with IL-1β, has been shown to increase tenocyte migration and proliferation, the capacity for which decreases with age.^80,83,87,88,100-106^ In this study, we observed a decrease in tenocyte migration and proliferation following inflammatory stimulation in all species (statistically significant for sheep tenocyte proliferation as well as rat and mouse tendon cell migration under constant inflammation) except rats (non-significant trend toward increased proliferation), which may be due to our use of tenocytes from individuals in disease-relevant age groups.

The overall gene expression of human tenocytes was most similar to murine under healthy, equine under transient and ovine under constant inflammatory conditions. The species difference between human and the four animal models was particularly evident in the expression of the main tendon matrix component COL1. Healthy tenocytes of all four model species exhibited significant differences to human in their expression of COL1. Col1 typically amounts to appr. 95% of total tendon collagen or 50-80% of tendon dry weight,^107^ but cytokines, such as IL-1β and TNF-α, suppress COL1 synthesis, which leads to reduced stiffness.^28,44,84,108^ In this study the decrease in COL1 synthesis following inflammatory stimulation could be observed in all species and was most pronounced in rat and mice, least in sheep and most similar to humans in horses.

The expression of the transcription factor SCX, a specific marker of the tendon/ligament lineage,^109^ while low in all species under healthy conditions, only increased in humans upon constant inflammation. SCX is a transcription factor that regulates tendon genes, including Col1 and Tnmd, and is required for normal tendon development^110,111^ and adult tendon repair in mice.^112,113^ An increase in its expression is likely to result in changes in the expression of its downstream genes and to be beneficial to tendon healing post injury.^113^ The essential contribution of SCX was also shown in SCX-null mice, which fail to convert from producing primarily COL3 to synthesizing mainly COL1 during tendon repair, supporting the hypothesis that the transcriptional control of collagen type I is mediated by SCX.^113^ Overall for the six tenogenic factors, rat tenocytes showed the largest difference to humans in the Mahalanobis distance, while tendon cells from mice and horses most closely equaled humans, indicating that these species might be most suitable for studies evaluating ECM production and tendon healing.

For matrix remodelling proteinases, the species differences were most prevalent for healthy tenocytes: tendon cells of all model species differed from humans for MMP13 and all but horses for MMP3. MMPs are key players in physiological and pathological tendon ECM remodeling, contributing to the degradation of tendon ECM and hence the loss of the biomechanical resistance and durability of tendon.^44,114-116^ An increase in MMP expression has also been implicated in the pathogenesis of tendinopathy.^44,116^ MMP13 specifically was upregulated in rotator cuff tendon tears and flexor tendon injury.^117-119^ In this study, inflammatory stimulation increased MMP13 expression in tenocytes of all species, only minimally in mice and horses but 4-8-fold in rats, sheep and humans. In contrast, all species showed similarly increased MMP1 expression following inflammatory stimulation; no significant differences were observed in MMP1 expression in any species in any condition compared to humans. This corresponds well with other studies showing upregulation of MMP1 in ruptured tendons suggesting a high level of collagen degradation by this enzyme.^120^ In total, for the functional group of ECM remodelling genes, horses again provided the best and rats the worst match to humans as shown in the Mahalanobis distance analysis.

For the expression of inflammatory mediators, the Mahalanobis distances of all species were larger than for the other functional gene groups. Although the immunophysiology of larger animal species has traditionally been presumed to be closer to humans than rodents,^47,121^ rat tenocytes most closely approximated human tendon cells in this category. Additionally, in healthy condition, mice presented the lowest distance from all animals, rising again the question if larger animals truly are more similar to human. Remarkably, healthy and transiently inflamed tenocytes of all four model species, as well as constantly inflamed ovine and equine tenocytes, showed significant differences to human in their expression of COX2. Following inflammatory stimulation, COX2 was only significantly upregulated in humans, mice and rats. Upregulation of COX2 plays an important, multifaceted role in the inflammatory cascade in injured tendons through the synthesis of prostaglandins.^125^ COX2 is essential in the early injury response as evidenced by impaired tendon repair following administration of selective COX2 inhibitors in the early repair phase.^122^ The lacking upregulation of sheep and horses therefore invites further investigation into the early tendon healing response in the different species *in vivo*.

Correspondingly, IL6, a cytokine with strong association with inflammation in tendon disease,^58,123,124^ displayed significantly different expression in transiently inflamed tenocytes of all species except rats. Statistically significant differences in IL6 expression compared to human were also evident under constant inflammatory conditions for mice and horses. IL6 plays an essential role in tendon healing as repair processes in IL6 knock-out mice are impaired.^38^ It tenocytes in two ways: i) IL6 stimulates tenocyte proliferation and survival and ii) it inhibits their tenogenic differentiation via the Janus tyrosine kinases/Stat3 signaling pathway.^44,125^

Cell properties may be influenced not only by species and interdonor differences but also by cell isolation and processing methods.^126,127^ In the present study, two isolation methods, enzymatic digestion and cell migration out of tendon explants, have been used depending on the available sample size. Enzymatic digestion was used for smaller sample sizes as higher cell yields are achieved with this method, while the explant technique is less invasive and requires less manipulation and labour. As both methods were used for all species and alterations in experimental conditions have been shown to be of minor importance to cell behaviour compared to cell source and interdonor variability,^126^ the isolation method is unlikely to have significantly influenced the species-specific gene expression profiles observed in this study.

In summary, the results of our study show that all four model species approximate some aspects of the behaviour of human tenocytes well and others poorly. No animal model sufficiently emulates human tenocytes’ cellular and molecular features and response to inflammation to be considered the gold-standard for tendon research. Translational medicine will need to continue to rely on a fit-for-purpose selection of animal models to approximate the human condition, based on the essential characteristics that must be mimicked for a particular research question.^19^ Peculiarities, strengths, and weaknesses of the model species need to be accounted for in the study design, analysis and interpretation.^19,128,129^ Data from multiple animal models should be combined to optimize translational predictive validity.

## Materials and Methods

Tenocytes of four mammalian species (mouse, rat, sheep, horse) were compared with human tenocytes (n = 3 donors, i.e., biological replicates, per species). All methods and experimental protocols in this study were carried out in accordance and compliance with relevant institutional and national guidelines and regulations.

### Tenocyte isolation from animals

All animals were euthanized for reasons unrelated to this study. Based on the “Good Scientific Practice. Ethics in Science und Research” regulation implemented at the University of Veterinary Medicine Vienna, the Institutional Ethics Committee (“Ethics and Animal Welfare Committee”) of the University of Veterinary Medicine Vienna does not require approval of in vitro cell culture studies, if the cells were isolated from tissue, which was obtained either solely for diagnostic or therapeutic purposes or in the course of institutionally and nationally approved experiments.

Species-specific, energy-storing, weight-bearing tendons were harvested from skeletally mature animals immediately following euthanasia: Achilles tendons from sheep (Merino-cross breed, female, aged 2-5 years), rats (Fischer344 breed, female, aged 3-4 months) and mice (c57bl/6 breed, female, aged 8-12 weeks); superficial digital flexor tendons from the front limb of horses (7-15 years, geldings). Under sterile conditions, the paratenon was removed and the tendons were sectioned into small pieces (<0.5 × 0.5 × 0.5 cm). Isolation of cells was performed either by enzymatic digestion using 3 mg/ml collagenase type II (Gibco Life technologies, Vienna, Austria) for 6-8 hours or migration from explants (explants were removed after 7-10 days) or a combination of both. Cells were expanded until 80 - 90% confluency before passaging.

### Human tenocytes

Human tenocytes obtained with ethical approval and informed consent from the Achilles tendon of three male human donors (aged 60-90 years) in accordance with relevant guidelines and regulations (Declaration of Helsinki) were purchased in cryopreserved condition in passage two from two different providers (Pelo Biotech GmbH, Germany and Zen-Bio, North Carolina, USA with review of the protocols and consent forms by an independent review board (Institutional Review Board, Pearl Pathways, LLC) which is accredited by the Association for the Accreditation of Human Research Protection Program Inc.).

### Cell culture

The culture medium was identical for all species: minimal essential medium (α-MEM, Sigma-Aldrich, Vienna, Austria) supplemented with 10% fetal bovine serum (FBS-12A, Capricon, Ebsdorfergrund, Germany), 1% L-Glutamine (L-Alanyl L-Glutamine 200Mm, Biochrom), 100 units mL^-1^penicillin and 0.1mg mL^-1^streptomycin (P/S, Sigma-Aldrich, Vienna, Austria). Cells were cultured at 37°C, 5% CO2 until the desired passage and number of cells was obtained. Experiments were performed with cells either in passage 3 or 4.

### Morphology

Cells were imaged both at low and high confluency using the EVOS FL Auto imaging system in phase contrast with a 40x and 400x objective (ThermoFisher Scientific, AMEP4680). Cell phenotypes and cell sheet patterns were characterised for all species and compared to human cells. Tenocyte dimensions (length and width) were measured for each of the five species.

### Inflammatory stimulation

Gene expression, proliferation and migration of tenocytes of all five species were compared under standard culture conditions (healthy control) as well as under transient (24 h) and constant exposure to inflammatory stimuli (10 ng/ml IL1β (Immuno Tools, Friesoythe, Germany) and 10 ng/ml TNFα (Immuno Tools, Friesoythe, Germany)).^Dakin:2018bi 96^ After 24-hour exposure to inflammation, the transient inflammation group received fresh culture medium, while for the constant inflammation group fresh medium was again supplemented with inflammatory factors (fig. 2).

### Proliferation assay

Tenocytes were plated in 96-well plates (3000 cells/well in technical triplicates) and cultured under control (healthy), transient and constant inflammatory conditions. The cell number per plate was quantified via DNA fluorescence using the CyQuant assay (Invitrogen) according to the manufacturer’s recommendations on day 0, 1, 2, and 3 (fig. 2). As cell proliferation sets in after a lag time of about 24 h and relative proliferation rates decrease steadily, we used log(cell nr) as the target variable and log(time in hours minus 23) as regression variable for the parametric statistical analysis.

### Wound healing (scratch) assay

Migration of tenocytes was evaluated in a wound healing model using a magnetic scratch device to create standardized cell-free gaps of 1.5 mm width in confluent sheets of tenocytes.^130^ Cells were seeded in 12-well plates (100,000 cells/well in technical triplicates) and left to adhere overnight. Inflammatory stimuli were added to the transient and constant inflammation groups and scratches were created 48 hours after seeding under control (healthy), transient and constant inflammatory conditions (fig. 2). The cell-free area was imaged at 24h intervals (0, 24, 48, 72, 96 hours, fig. 2) in phase contrast using the EVOS FL Auto imaging system with a 4x fluorite objective using coordinate recovery function. The gap size was measured using the MRI Wound healing Tool (http://dev.mri.cnrs.fr/projects/imagej-macros/wiki/Wound_Healing_Tool) in ImageJ (https://imagej.nih.gov/ij/, version 2.0.0-rc-43/1.50e). As gap closure approached 100% in the fastest group, healthy rat tenocytes, at 48h, this time point was chosen as cut-off for slope calculations and comparison of conditions. For the parametric analysis, we used the untransformed gap area [mm^2^] as target variable and the untransformed time between 0 hours and 48 hours (before the gap closed in any of the samples) as regression variable.

### Quantitative PCR

Tenocytes were seeded in 12-well plates (100,000 cells/well in technical triplicates) and cultured under control (healthy), transient and constant inflammatory conditions. Cells were harvested for RNA isolation using RNA isolation reagent (Trizol, ThermoFisher Scientific, MA, USA) 48 hours after initiation of inflammation, as previously described.^131^ The 48 hours time point was chosen as it allows assessment of the response to inflammation as well as to removal of inflammatory stimuli.

Briefly, a solution of Trizol and Chloroform (Sigma-Aldrich) in a ratio of 5 to 1 was used. Total RNA was recovered by the addition of isopropyl alcohol (Sigma-Aldrich) and glycerol (Thermo Scientific). The mixture was incubated on ice and centrifuged for 45 minutes at 13,000 rpm. The total RNA pellet was washed with 75% ethanol and solubilized in RNase-free water. Genomic DNA was removed by a DNA removal kit (Life Technologies, Carlsbad, California, USA). Two nanograms of RNA from each sample was used for the qPCR reaction (qPCR One-Step Eva Green kit, Bio&Sell, Feucht, Germany).

We measured gene expression of tendon markers (TNC, TNDM, SCX), collagens (COL1, COL3, COL5), matrix-metalloproteinases (MMP1, MMP3, MMP13), inflammatory factors (IL6, COX2, NFkB, p53), a marker for aberrant tenocyte differentiation (ALP) and for focal adhesion and migration (FAK) in the four model species and humans under healthy, transiently and constantly inflamed conditions. All primers were designed using the Primer3 software. Primer sequences are shown in supplementary table 2. The transcript level for the 15 genes of interest was normalized to the transcript level of the housekeeping gene glyceraldehyde-3-phosphate dehydrogenase (GAPDH) and presented as ratio to GAPDH.^58^ The ratio between COL 1 and COL 3 was also evaluated for further matrix remodelling characterization, with a higher COL1:COL3 ratio indicating a stronger tenogenic phenotype.^132,133^ For the parametric analysis, we used the log_2_ transformed ratios of the target gene to GAPDH as target variable.

### Statistical analysis

For statistical analyses, the R statistical programming language^134^ and GraphPad (version 8.4.2) were used. Target and regression variables (where appropriate) are given in the respective subsections. Data are presented descriptively as mean and standard deviation. Generally, linear models (analyses of variance and covariances, ANCOVA) were used, e.g., for wound healing the untransformed area was the target variable, *time* a regression variable, and *species* and *condition* factors; as interactions of two explanatory variables *time*species, time*condition, condition*species*, and *biological replicate* nested within *species* were included; furthermore, the three-way interaction *time*species*condition* was also included. Note that all terms with *time* are to be interpreted as slopes or differences in slopes. The Tukey’s HSD (honestly significant difference) test was used to account for multiple testing, where appropriate. Confidence intervals of parameter estimates were calculated.

Note that many different target variables are available, i.e., data are multidimensional. With qPCR alone, 15 genes of interest were measured under three conditions. For each gene separately, an ANOVA with species and condition was calculated. Furthermore, we condensed information by calculating the multivariate (Mahalanobis) distance of the log_2_-transformed mRNA concentrations, for the three conditions of each gene, and report the distance of each of the four mammalian species from the human values. For a single condition, all 15 genes of interest could be used for calculating the multivariate distance. Additionally, we grouped genes into classes, e.g., all collagens or all matrix-metalloproteinases and calculated multivariate distances for the classes separately, this time jointly for the different conditions. We also calculated a principal component analysis (PCA) of the log_2_-transformed qPCR data for all gene, treatment, and species combinations together. The proportions of the variance explained by the different PC’s are reported and the rotated data for the different treatment and species combinations are shown in graphs for the most important components.

## Supporting information

Supplementary figure 1: Graph of the Mahalonobis distances of the four model species to human for each gene

Supplementary figure 2: Pairwise plots of a Principal Component Analysis of gene expression values with PC1 explaining 36% of the variance, PC2 23%, P

Supplementary table 1: The qPCR results (log2 FC relative to GAPDH) for each of the 15 genes (3 biological replicates/species) are listed for each con

Supplementary table 2: qPCR Primer sequences

## Acknowledgements

The authors acknowledge Sinan Gültekin for his technical and John Breteler for his graphical support. This research was supported by the Austrian Research Promotion agency (grant number 7269695) and the University of Vienna tandem PhD programme.

