## Supplementary figures and images for "Species variations in tenocytes’ response to inflammation require careful selection of animal models for tendon research"

### Supplementary figure 1: Graph of the Mahalonobis distances of the four model species to human for each gene

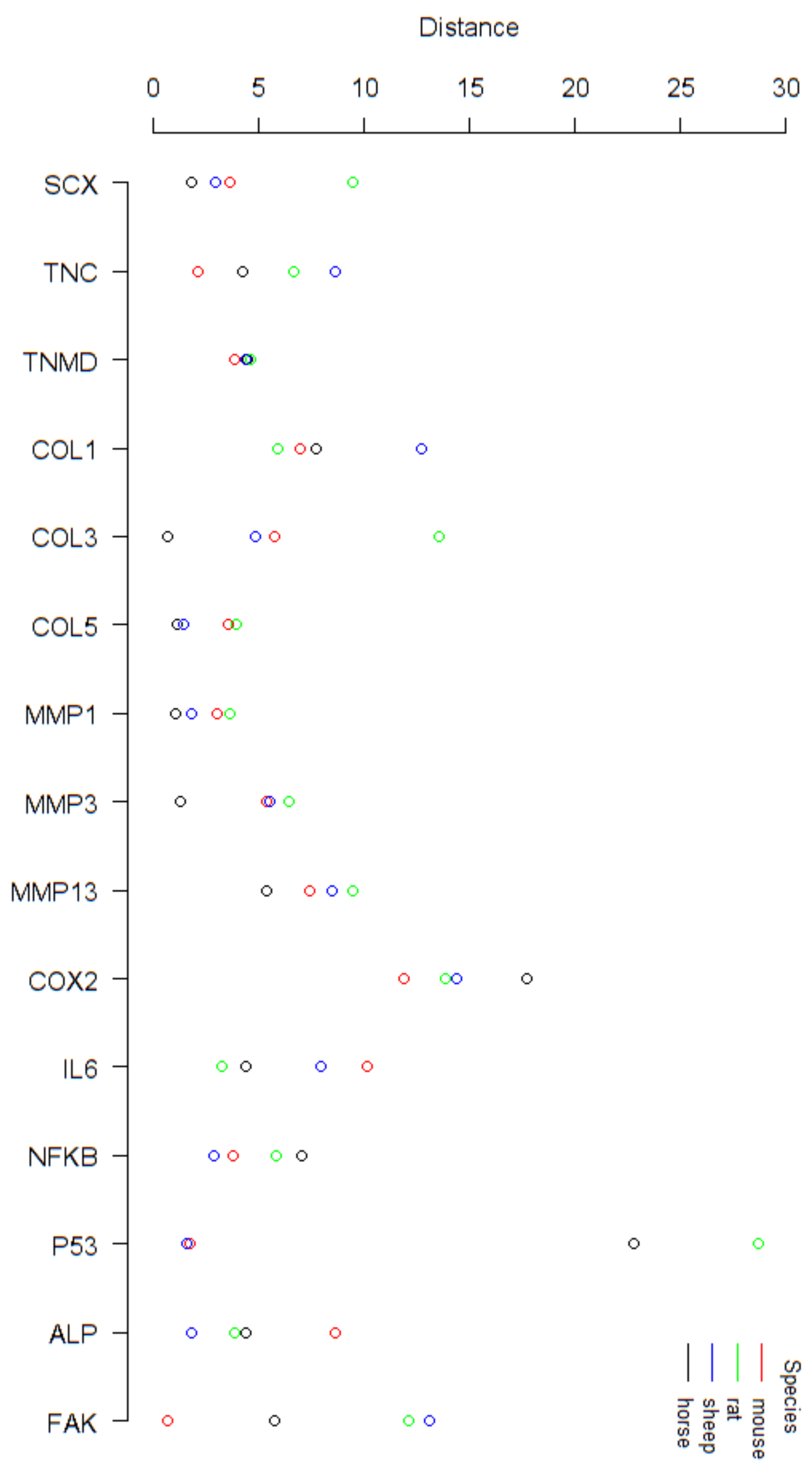
