## Supplementary figure 2: Pairwise plots of a Principal Component Analysis of gene expression values with PC1 explaining 36% of the variance, PC2 23%, P for "Species variations in tenocytes’ response to inflammation require careful selection of animal models for tendon research"

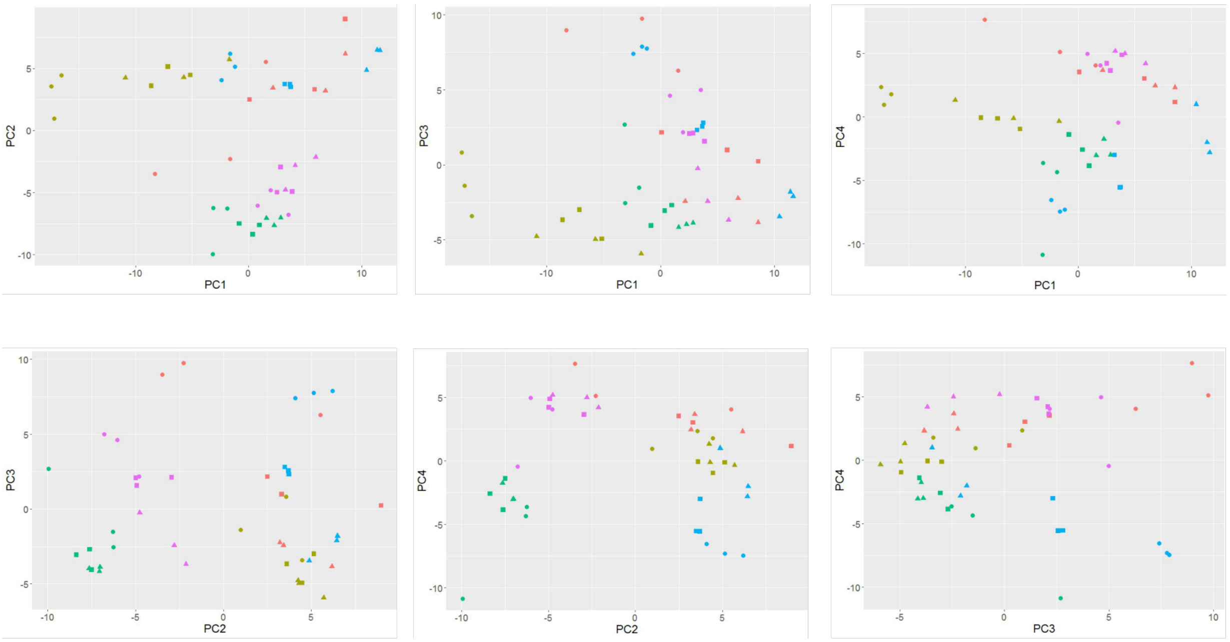


Supplementary figure 2: Pairwise plots of a Principal Component Analysis of gene expression values with PC1 explaining 36% of the variance, PC2 23%, PC3 14% and PC4 13%. Species are colour coded, conditions (healthy, transiently inflamed and continuously inflamed) are differentiated by symbols.
