## Supplementary table 1: The qPCR results (log2 FC relative to GAPDH) for each of the 15 genes (3 biological replicates/species) are listed for each con for "Species variations in tenocytes’ response to inflammation require careful selection of animal models for tendon research"

| | HEALTHY | | | | | TRANSIENT INFLAMMATION | | | | | CONSTANT INFLAMMATION | | | | |
|---|---|---|---|---|---|---|---|---|---|---|---|---|---|---|---|
| | mouse | rat | sheep | horse | human | mouse | rat | sheep | horse | human | mouse | rat | sheep | horse | human |
| SCX | -5.608 | -2.351 | -3.336 | -5.000 | -7.158 | 0.000 | -1.445 | -0.500 | 0.760 | 0.837 | 0.597 | -1.131 | -3.629 | -2.178 | 0.280 |
| | -3.718 | -1.905 | -8.158 | -5.484 | -8.158 | -1.119 | -2.333 | 1.716 | -1.474 | 0.515 | -1.248 | -1.911 | 2.837 | -2.996 | 0.893 |
| | -5.184 | -2.377 | -3.816 | -7.520 | -7.158 | -0.196 | -2.706 | -0.626 | 0.759 | 1.000 | -0.138 | -3.888 | -2.342 | -1.231 | 1.716 |
| TNC | -3.191 | -0.029 | 3.410 | -2.260 | -2.012 | -0.243 | -0.522 | -1.522 | 2.341 | 0.890 | -0.536 | -1.331 | -1.365 | 3.016 | 0.406 |
| | -2.751 | 0.561 | 0.973 | -2.885 | -2.680 | -0.307 | -0.845 | 0.914 | 0.656 | -0.023 | -0.549 | -1.467 | 0.058 | 0.389 | -0.465 |
| | -3.191 | 1.103 | 0.100 | -2.465 | -2.208 | -0.307 | -1.700 | 1.202 | 2.538 | 0.335 | -0.408 | -2.445 | -0.536 | 3.470 | 0.097 |
| TNMD | -4.540 | -6.108 | -8.040 | -6.012 | -6.265 | -2.178 | -1.536 | 2.511 | 0.090 | 2.000 | -1.671 | -1.688 | 2.140 | -0.867 | 3.862 |
| | -3.943 | -5.718 | -4.150 | -6.506 | -2.816 | -1.774 | -1.341 | 0.200 | 0.585 | 0.638 | -2.115 | -2.663 | -0.905 | -0.759 | 0.853 |
| | -5.238 | -2.873 | -8.966 | -7.381 | -1.500 | -1.728 | -3.569 | 1.170 | 0.415 | -0.535 | -1.728 | -5.771 | -1.000 | 0.874 | -0.249 |
| COL1 | -1.355 | 5.466 | -4.958 | -1.000 | 2.013 | -1.918 | -2.209 | -2.696 | 0.711 | -0.043 | -2.461 | -3.858 | -1.792 | -1.295 | -1.352 |
| | 0.330 | 5.803 | -6.005 | -3.065 | 1.804 | -4.156 | -2.773 | -0.280 | 0.927 | -0.380 | -4.696 | -4.588 | -0.959 | 0.153 | -1.079 |
| | -2.252 | 6.384 | -6.966 | -4.294 | 1.987 | -2.485 | -4.523 | 1.907 | 1.839 | -0.289 | -2.627 | -7.089 | 0.322 | 1.139 | -1.690 |
| COL3 | 1.654 | 2.929 | 1.371 | 0.990 | -0.724 | -1.283 | -2.571 | -2.357 | 1.520 | 1.699 | -1.486 | -5.034 | -1.521 | 0.380 | 1.960 |
| | 2.575 | 3.054 | -1.785 | -0.629 | -0.806 | -1.794 | -2.701 | -0.184 | 0.406 | 1.399 | -2.620 | -5.536 | -0.215 | 0.885 | 1.196 |
| | 1.537 | 3.557 | -2.932 | -0.975 | -0.316 | -1.731 | -3.640 | 1.395 | 1.386 | 1.285 | -2.244 | -7.363 | -0.145 | 1.111 | 0.394 |
| COL5 | -7.966 | 1.862 | -0.987 | -0.118 | -4.351 | -2.000 | -1.829 | -1.811 | 0.504 | 2.215 | 1.907 | -2.819 | -2.128 | -0.900 | 4.393 |
| | -1.730 | 2.577 | -3.020 | -0.143 | -1.749 | -2.329 | -2.491 | 0.146 | -0.391 | 1.177 | -2.988 | -3.669 | -0.469 | -0.970 | 1.340 |
| | -3.224 | 3.016 | -1.163 | -1.015 | -0.304 | -1.187 | -2.692 | 0.903 | 0.728 | 0.312 | -2.127 | -5.007 | -1.860 | 0.120 | -1.372 |
| MMP1 | -10.85 | -7.966 | -11.57 | -1.518 | -6.442 | 0.852 | 1.392 | 4.995 | 4.133 | 5.063 | 1.228 | 8.047 | 8.630 | 3.501 | 6.242 |
| | -20.41 | -7.796 | -9.409 | -10.67 | -2.506 | 9.211 | 1.152 | 2.213 | 7.661 | -1.202 | 11.13 | 8.124 | 5.061 | 8.264 | 0.217 |
| | -11.25 | -7.644 | -7.644 | -11.08 | -5.540 | 0.450 | 2.293 | 0.766 | 6.934 | 2.953 | 2.311 | 7.837 | 0.585 | 8.758 | 3.295 |
| MMP3 | -0.135 | 0.647 | 2.098 | -2.601 | -8.966 | 0.540 | 3.123 | -3.000 | 3.151 | 6.618 | 1.878 | 5.292 | -0.880 | 3.138 | 7.916 |
| | 1.069 | 1.759 | -1.026 | -7.998 | -8.381 | 0.105 | 2.362 | 0.633 | 6.393 | 3.544 | 0.806 | 4.119 | 0.230 | 7.795 | 4.992 |
| | 0.112 | 0.991 | -1.038 | -10.90 | -9.966 | 0.879 | 1.631 | -0.477 | 6.269 | 6.615 | 1.354 | 3.200 | -1.045 | 6.852 | 5.585 |
| MMP13 | -2.868 | -5.608 | 0.088 | -3.980 | -10.97 | -0.639 | 3.239 | -0.575 | 4.820 | 6.755 | 0.859 | 7.735 | 0.037 | 5.800 | 7.358 |
| | -1.519 | -5.718 | -1.504 | -2.990 | -10.97 | -0.323 | 3.359 | 0.173 | 4.719 | 3.459 | 0.622 | 7.741 | 1.142 | 4.821 | 4.585 |
| | -2.006 | -4.837 | -2.065 | -8.820 | -9.966 | -0.616 | 2.010 | 1.138 | 4.200 | 3.087 | 0.225 | 6.257 | 1.987 | 5.800 | 0.000 |
| COX2 | -3.837 | -2.366 | 0.600 | 3.341 | -13.80 | 2.396 | 2.525 | 0.898 | 0.662 | 7.158 | 3.621 | 6.687 | 0.612 | -0.536 | 11.05 |
| | -4.308 | -1.706 | 1.066 | 3.491 | -13.29 | 3.557 | 1.389 | 0.389 | -0.306 | 7.180 | 4.450 | 5.719 | -0.430 | -1.341 | 8.607 |
| | -3.458 | -1.831 | -0.624 | 0.335 | -12.70 | 2.673 | 2.638 | 1.182 | 0.827 | 6.737 | 3.758 | 6.168 | 0.484 | 0.506 | 4.059 |
| IL6 | -0.748 | -4.680 | -4.426 | -7.758 | -4.573 | 1.131 | 4.387 | 1.796 | 8.568 | 4.795 | 2.137 | 8.027 | 6.273 | 11.15 | 6.773 |
| | 0.759 | -4.012 | -7.059 | -8.105 | -3.591 | 0.572 | 4.100 | 2.679 | 7.525 | 3.580 | 0.902 | 6.470 | 7.847 | 10.68 | 5.252 |
| | -0.276 | -3.911 | -3.211 | -9.380 | -6.059 | 0.526 | 2.858 | 1.325 | 7.956 | 5.924 | 1.164 | 6.728 | 2.458 | 12.58 | 5.805 |
| NFKB | -6.506 | -3.826 | -7.644 | -0.201 | -5.878 | -0.212 | 1.671 | 1.070 | 0.331 | 1.122 | 0.241 | 1.966 | 0.848 | -1.187 | 2.031 |
| | -5.108 | -2.552 | -6.573 | -1.857 | -5.837 | -0.904 | 0.338 | 0.308 | 0.391 | 0.456 | -0.951 | 0.913 | 1.280 | 0.262 | 1.314 |
| | -6.211 | -1.855 | -5.158 | -3.786 | -5.506 | -0.433 | 0.122 | -0.763 | 0.955 | 0.585 | -0.755 | 0.049 | -0.135 | 1.558 | -0.816 |
| P53 | -5.411 | -1.095 | -5.059 | -1.042 | -5.351 | -1.095 | -0.519 | -1.000 | 0.894 | -0.708 | -1.031 | -0.039 | -1.447 | -0.912 | -0.661 |
| | -5.059 | -0.931 | -5.644 | -1.521 | -5.351 | -1.737 | -0.158 | -0.515 | 0.069 | -0.405 | -1.447 | -0.069 | -1.000 | -0.243 | -0.615 |
| | -6.211 | -0.336 | -6.158 | -2.671 | -5.083 | -0.948 | -2.339 | 0.000 | 1.191 | -0.883 | -0.848 | -2.415 | -0.052 | 0.925 | -1.795 |
| ALP | -4.071 | -3.184 | -5.878 | -5.680 | -4.921 | -1.725 | -2.026 | 0.795 | 1.972 | 0.222 | 2.136 | -2.026 | 0.957 | 2.063 | -0.835 |
| | -1.236 | -3.573 | -5.322 | -6.265 | -3.955 | 0.836 | -2.222 | 2.287 | -0.177 | -1.367 | 1.332 | -2.144 | -0.515 | -0.379 | -0.924 |
| | -0.853 | -3.272 | -4.816 | -6.966 | -6.718 | 0.323 | -2.301 | -0.864 | 2.919 | -0.788 | 0.606 | -3.524 | -1.150 | 3.426 | -2.663 |
| FAK | -7.172 | -2.454 | -4.265 | -4.055 | -7.265 | -0.191 | -1.501 | 2.611 | 0.889 | -0.115 | -1.345 | -1.501 | 2.095 | -0.922 | -1.700 |
| | -7.199 | -2.035 | -1.493 | -4.495 | -7.265 | -0.427 | -1.761 | -0.642 | 0.277 | -0.700 | -0.435 | -2.316 | -0.922 | -0.155 | -0.115 |
| | -7.012 | -1.991 | -3.041 | -5.744 | -6.108 | -1.860 | -2.274 | 0.876 | 0.674 | -0.273 | -0.535 | -3.220 | 0.299 | -0.171 | -1.158 |
