## Supplementary table 2: qPCR Primer sequences for "Species variations in tenocytes’ response to inflammation require careful selection of animal models for tendon research"

| | mouse | rat | sheep | horse | human |
|---|---|---|---|---|---|
| **COL 1** | | | | | |
| F 5'-3' | GTGCTAAGGGTGAAGCTGG | CGGCAGAAGTCTCAAGATGGTGGCCG | AGTTGGTCGAACTGGAGAGC | TCCATCTGGAGAGCCTGGTA | ACATGTTCAGCTTTGTGGACCTCCG |
| R 5'-3' | CATCAGCACCAGGGTTTCCAG | CTCTCCGCTCTTCCAGTCAGA | GAGACCCAGAAAACCAGGAG | CACCTGGTAGACCACGTTCA | ACGCAGGTGATTGGTGGGATGTCT |
| **COL 3** | | | | | |
| F 5'-3' | CTGGAGATAAGGGTGAAGGT | TCCCCTGGAATCTGTGAATC | AAGGATGGGACAAGTGGACA | TGGAGCTCCTGGACTGATAG | CGGCAATCCTGAACTTCCTG |
| R 5'-3' | GAGGGCCTCCTTCACCTTTCT | CCGACTCCAGACTTGACATC | GGAGCCCTCAGATCCTCTTT | CCATTCTTACCAGGCTCACC | ATCAGCTTCAGGGCCTTCTT |
| **COL 5** | | | | | |
| F 5'-3' | CAGTGAATTCAAGCGTGGGA | GGATGTTGCCTACCGAGTCT | TGTTTGAGGGTGACATCCAG | TAGACGTCAACGGCATCATC | GGCATCGGGGACTATGACTA |
| R 5'-3' | GTAGGTGACGTTCTGGTGGG | CTGCCTTTCTTGGCTTTCAC | TCCTGAGACTGGGGTTTGTC | TGGTCTGAGACGAAGAGCAG | CTCGCCATAGGTGAGGTCAT |
| **MMP 1** | | | | | |
| F 5'-3' | TATGGACCTTCCCCAAATCC | GCCCATACAGTTTGAATACAGTATCTG | GACCTGGTGGAAACCTTGCT | CAGTGCCTTCAGAAACACGA | GCAGCTTCAGTGACAAAC |
| R 5'-3' | CCGAATGTAGAACCTGCCTTT | CCAGTTTAATAAACACCATCTCTTGA | AGCTGCCACCCGATACAAGT | GCTTCCCAGTCACTTTCAGC | TTCCTCCGGCAAAGACTCAT |
| **MMP 3** | | | | | |
| F 5'-3' | CCTATTCCTGGTTGCTGCTC | TGGACCAGGGATTAATGGAG | TGTGATCCTGCCTTGTCCT | TGTGGAGGTGATGCACAAATC | GTAGAAGGCACAATATGG |
| R 5'-3' | AACGGGACAAGTCTGTGGAG | GAGCAGCAACCAGGAATAGG | ATGCCTGAAGGAAGAGATGG | GCATGCCAGGAAATGTAGTGAA | ACTCTATGTGACAAGGTG |
| **MMP 13** | | | | | |
| F 5'-3' | CTTTTCCTCCTGGACCAAACT | CCCTCGAACACTCAAATGGT | GGTCTGTTGGCTCACGCTTTCC | TGGTCCAGGAGATGAAGACC | TCAGGAAACCAGGTCTGGAG |
| R 5'-3' | TCATGGGCAGCAACAATAAA | AAGGCCTTCTCCACTTCAGA | GAGTGCTCCTGGGTCCTTGG | GATGGCATCAAGGGATAAGG | TGACGCGAACAATACGGTTA |
| **IL 6** | | | | | |
| F 5'-3' | GTTCTCTGGGAAATCGTGGA | TGTATGAACAGCGATGATG | GTCATGGAGTTGCAGAGCAG | ATGGCAGAAAAAGACGGATG | CTCAGCCCTGAGAAAGGAGA |
| R 5'-3' | TTCTGCAAGTGCATCATCGT | AGAAGACCAGAGCAGATT | ACCCACTCGTTTGAGGACTG | GGGTCAGGGGTGGTTACTTC | AGGTTGTTTTCTGCCAGTGC |
| **COX 2** | | | | | |
| F 5'-3' | TGACAGTCCACCTACTTACAAT | GCAAATCCTTGCTGTTCCAACCCA | CCGAAGATGTCCTACCCTCA | CGAGTGGTTCTCCCCATAGA | TTCAAATGAGATTGTGGGAAAAT |
| R 5'-3' | CTCCACCAATGACCTGATAT | TTGGGGATCCGGGATGAACTCTCT | CGTAGAATAGGCCTGGACGA | GGCCACGAGAGTTGTCTGAT | AGATCATCTCTGCCTGAGTATCTT |
| **SCX** | | | | | |
| F 5'-3' | TTGAGCAAAGACCGTGACAGA | AGGGCCTGTGAACAGAGAGA | TCTGCCTCAGCAACCAGAGA | TCTGCCTCAGCAACCAGAGA | CAGATCTGCACCTTCTGCCT |
| R 5'-3' | TGTGGACCCTCCTCCTTCTAAC | AGGTAGAGAGCCAGCATGGA | TCCGAATCGCCGTCTTTC | TCCGAATCGCCGTCTTTC | GAATCGCTGTCTTTCTGTCGC |
| **TNMD** | | | | | |
| F 5'-3' | GTCACATTCTAAATGCAGAAG | AAGACCTATGGCATGGAGCACA | TGGGGGAGTAAGCACTTCTG | GGCATCTACTTCGTGGGTCT | GGCATCTACTTCGTGGGTCT |
| R 5'-3' | CTCCCCCAAAACAGGACAAT | CGGATCAAAGTAGATGCCAGTGTATCCG | CAGTGCCATTTCCACTTCTG | TTCTGCTGGGACCCAAATCA | TTCTGCTGGGACCCAAATCA |
| **TNC** | | | | | |
| F 5'-3' | CTGCTGTCAAGGGAGACAAG | GCCAGAGTTGCCACCTACTT | GCCATCCTGGAGAACAAGAA | ACCTCAGAGAAGGGCAGACA | CCAGCCAAAGAGACCTTCAC |
| R 5'-3' | AGACACCCGTAAGTCCTTGG | ATGTCCAGAGGATCCCACTC | ATCCCACTCCACTTCCACAG | CACCGTGAGGTTTTCCAGTT | TCCAGGGTGATGCTGTTATC |
| **GAPDH** | | | | | |
| F 5'-3' | CTGCACCACCAACTGCTTAG | GGCACAGTCAAGGCTGAGAATG | AGATGGTGAAGGTCGGAGTG | GTTTGTGATGGGCGTGAAC | ATCCCATCACCATCTTCCAG |
| R 5'-3' | GTCTTCTGGGTGGCAGTGAT | ATGGTGGTGAAGACGCCAGTA | GAAGGTCAATGAAGGGGTCA | GATGCCAAAGTGGTCATGG | TGGACTCCACGACGTACTCA |
